# An mRNA vaccine candidate for the SARS-CoV-2 Omicron variant

**DOI:** 10.1101/2022.02.07.479348

**Authors:** Jinkai Zang, Chao Zhang, Yannan Yin, Shiqi Xu, Weihua Qiao, Dimitri Lavillette, Haikun Wang, Zhong Huang

## Abstract

The newly emerged Omicron variant of severe acute respiratory syndrome coronavirus 2 (SARS-CoV-2) contains more than 30 mutations on the spike protein, 15 of which are located within the receptor binding domain (RBD). Consequently, Omicron is able to extensively escape existing neutralizing antibodies and may therefore compromise the efficacy of current vaccines based on the original strain, highlighting the importance and urgency of developing effective vaccines against Omicron. Here we report the rapid generation and evaluation of an mRNA vaccine candidate specific to Omicron. This mRNA vaccine encodes the RBD of Omicron (designated RBD-O) and is formulated with lipid nanoparticle. Two doses of the RBD-O mRNA vaccine efficiently induce neutralizing antibodies in mice; however, the antisera are effective only on the Omicron variant but not on the wildtype and Delta strains, indicating a narrow neutralization spectrum. It is noted that the neutralization profile of the RBD-O mRNA vaccine is opposite to that observed for the mRNA vaccine expressing the wildtype RBD (RBD-WT). Our work demonstrates the feasibility and potency of an RBD-based mRNA vaccine specific to Omicron, providing important information for further development of bivalent or multivalent SARS-CoV-2 vaccines with broad-spectrum efficacy.

## Main text

The newly emerged Omicron (B.1.1.529) variant of severe acute respiratory syndrome coronavirus 2 (SARS-CoV-2) was declared a variant of concern (VOC) by the World Health Organization (WHO) on 26^th^ November, 2021, and since then has spread globally at an unprecedented speed^1,2^. Unlike other VOCs, Omicron harbors a large number of mutations, including more than 30 mutations on the spike protein, 15 of which are located within the receptor binding domain (RBD)^3^. As a consequence, Omicron gains the ability to extensively escape vaccine- and infection-elicited neutralizing antibodies in vitro ^4,5–7^, raising a serious concern about the efficacy of existing vaccines in a setting where Omicron circulates predominantly. It is therefore of significant importance to develop new vaccine candidates targeting the Omicron variant as a part of the preparedness plan.

In this study, we adopted a non-replicating mRNA vaccine platform to rapidly produce and evaluate candidate vaccines against the Omicron variant. Two mRNA constructs encoding the RBD from the SARS-CoV-2 wildtype (WT) strain or from the Omicron variant were generated, designated RBD-WT and RBD-O, respectively (Fig. 1A). To verify the constructs, HEK293T cells were transfected with *in vitro* transcribed mRNA and subsequently analyzed for expression of the target proteins. Both RBD-WT and RBD-O mRNA-transfected cells, but not the control cells transfected with the luciferase-expressing mRNA, could be positively detected by an anti-RBD polyclonal antibody in the immunofluorescence staining assays (Fig. 1B). For both RBD-WT and RBD-O mRNA-transfected samples, the target proteins (~35 KDa, which is close to the predicated molecular mass) could be detected in the culture medium by both anti-His-tag and anti-RBD antibodies, indicating that the target proteins were correctly expressed and secreted (Fig. 1C). Together, the above results validate the mRNA vaccine constructs.

**Fig. 1.**
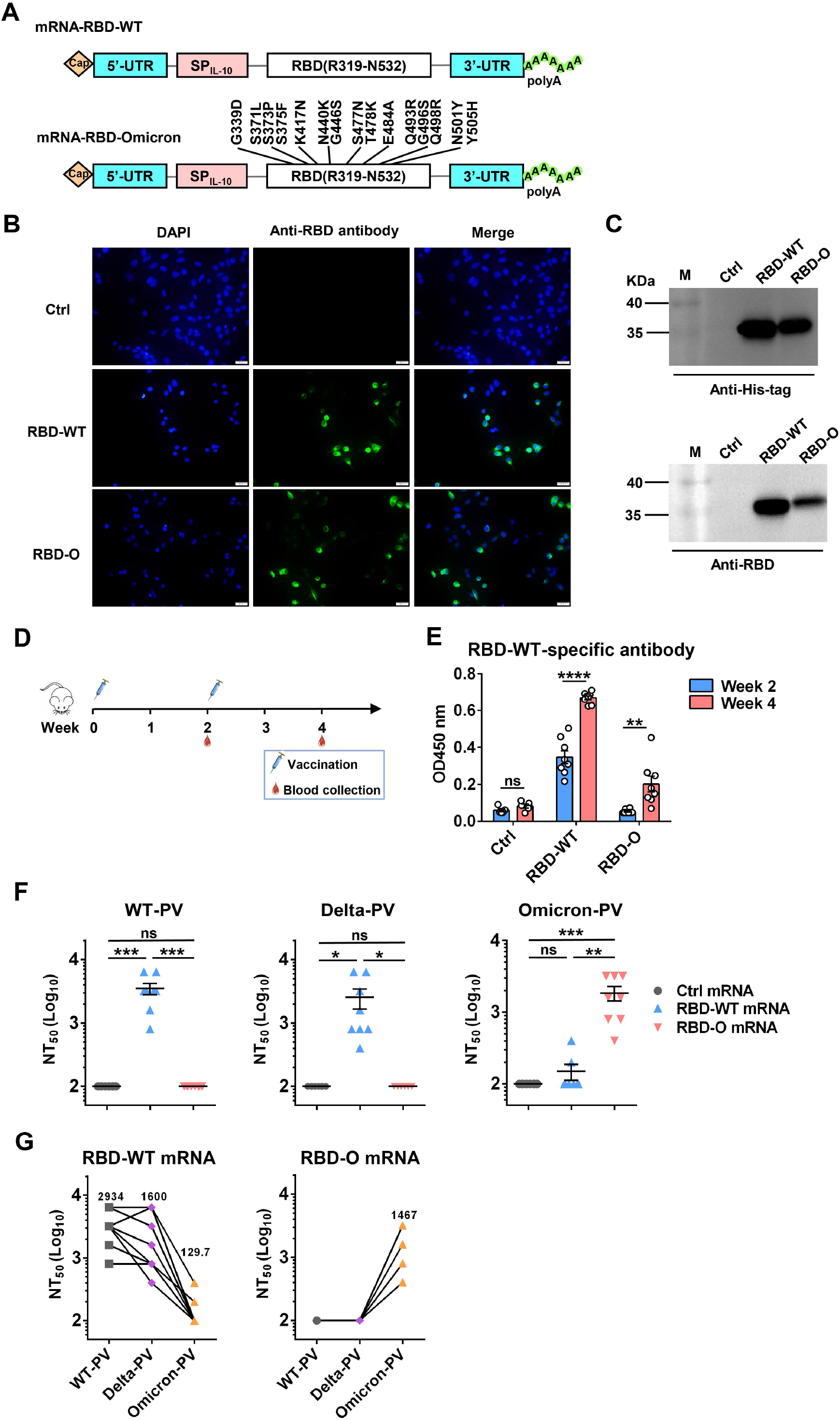
Design, characterization and immunogenicity assessment of LNP-encapsulated RBD mRNA vaccines against SARS-CoV-2. **(A)** Schematic diagram of RBD-WT and RBD-Omicron mRNA framework. Note that 15 mutations are located within Omicron RBD. UTR, untranslated regions; SP_IL-10_, human interleukin-10 signal peptide. **(B)** *In vitro* transcribed mRNA was transfected into HEK293T cells and the expression of RBD proteins within the cells were analyzed by immunofluorescence staining analysis with mouse anti-RBD polyclonal antibody and anti-mouse Alexa Fluor^®^ 488 secondary antibody. Ctrl, mRNA encoding luciferase. RBD-O, mRNA encoding RBD-Omicron. Overlay, merge of the blue (DAPI) and green (RBD) channels. Scale bars = 20 μm. **(C)** The culture supernatants of mRNA-transfected HEK293T cells were analyzed for RBD expression by western blotting with HRP-conjugated anti-His tag antibody and anti-RBD polyclonal antibody as detection antibodies. M, marker. **(D)** Mice immunization schedule. Groups of BALB/c mice were injected i.m. with 10 μg of RBD-WT mRNA (n = 8), RBD-O mRNA (n = 8), or luciferase-mRNA (Ctrl; n = 5) vaccines at weeks 0 and 2. Serum samples were collected from individual mice at weeks 2 and 4. **(E)** The week-2 and week-4 antisera were diluted 1:100 and analyzed for RBD-WT-binding activity by ELISA. **(F)** Neutralizing titers of the week-4 antisera samples from each group against WT-, Delta-, and Omicron-PV (pseudovirus). Serum samples that exhibited less than 50% neutralization at the lowest serum dilution (1:200) were assigned a NT_50_ value of 100 for statistical analysis. Each symbol represents one mouse. **(G)** Neutralizing titers of the week-4 antisera from RBD-WT mRNA group (left panel) and RBD-O mRNA group (right panel) against SARS-CoV-2 PV. The geometric mean of NT_50_ values was shown. For panels **E-F**, data are presented as mean ± SEM. *p* values were analyzed with unpaired *t*-test and indicated as follows: ns, not significant; *, *p* < 0.05; **, *p* < 0.01; ***, *p* < 0.001; ****, *p* < 0.0001.

Next, we encapsulated the antigen-expressing mRNAs with lipid nanoparticles (LNPs) to generate mRNA vaccine stocks. To visualize the in vivo delivery and expression of the mRNA vaccines, a firefly luciferase (FLuc) reporter-encoding mRNA-LNP formulation prepared using the identical methods was injected into mice, followed by bioluminescence imaging at different time points. Strong bioluminescence was observed in the FLuc mRNA-LNP injected mice at 6 hours post injection, indicating robust expression of FLuc (Supplementary Fig. S1). The bioluminescent signal then decreased gradually but remained detectable up to 48 hours post injection in some mice (Supplementary Fig. S1). These results demonstrate that our mRNA-LNP formulations are capable of delivering mRNA into cells and expressing the target antigens in vivo.

We then performed mouse immunization experiments to assess the immunogenicity and efficacy of the mRNA vaccine candidates. Two groups of female BALB/c mice (8 animals/group) were intramuscularly (i.m.) immunized with the RBD-WT or the RBD-O mRNA vaccine, respectively, at weeks 0 and 2 (Fig. 1D). Another group of mice (5 animals/group) were injected with the FLuc-expressing mRNA-LNP formulation, serving as the control in the experiment. Serum samples were collected from individual mice at weeks 2 and 4 for antibody titration and functional analysis (Fig. 1D).

The antisera were first evaluated for the presence of antigen-specific antibodies by ELISA with mammalian cell-produced recombinant RBD-WT as the capture antigen. All antisera were diluted 1:100 and then subjected to ELISA. The results were shown in Fig. 1E. The sera from the control (Fluc mRNA) group produced only background level of OD450 nm reading. In contrast, strong RBD-WT binding activity was observed for the RBD-WT mRNA vaccine sera collected at week-2 and the binding signal increased significantly at week-4, demonstrating the elicitation of antigen-specific binding antibodies by the mRNA vaccine. In addition, none of the week-2 antisera from the RBD-O mRNA vaccine group exhibited significant binding activity to RBD-WT, however, low levels of RBD-WT binding were detected in some of the week-4 antisera, suggesting that the RBD-O mRNA vaccine is immunogenic despite its ability to induce cross-reactive antibodies is limited. We also produced mammalian cell-expressed recombinant Omicron RBD protein. However, this protein exhibited aberrant binding orientation/coating efficiency on ELISA plate (data not shown) probably due to its surface property change resulting from the large number of amino acid substitution, and therefore was not used as the coating antigen in ELISA-based antibody measurement.

The week-4 antisera were assessed for their neutralization potency and breadth against a small panel of SARS-CoV-2 pseudovirus (PV), including the WT-PV, Delta-PV, and Omicron-PV (Fig. 1F). As expected, none of the antisera from the control (FLuc) group displayed neutralization towards any of the three PVs. Note that the serum samples unable to produce ≥50% neutralization when diluted 1:200 (the lowest serum dilution tested in the neutralization assays) were assigned a titer of 100 for geometric mean titer (GMT) computation. In contrast, all antisera from the RBD-WT mRNA vaccine group could potently neutralize the WT-PV with titers ranging from 800 to 6400 (GMT=2934) and also neutralize the Delta-PV with a slightly decreased efficiency (GMT=1600), however, only two out of the eight antisera marginally neutralized the Omicron-PV with titers being 200 and 400, respectively. For the RBD-O mRNA vaccine group, all antisera efficiently neutralized the Omicron-PV with a neutralizing GMT of 1467, but none of them exhibited cross-neutralization against the WT-PV and Delta-PV (Fig. 1F).

To better characterize the antisera’s cross-neutralization capacity, the NT50 titers of individual mouse sera against the three PVs were directly compared (Fig. 1G). For individual RBD-WT mRNA vaccine sera, their neutralization potency appeared not significantly affected by the Delta variant (less than 2-fold change in neutralizing GMT) but decreased drastically or even completely lost towards the Omicron variant. Conversely, all of the RBD-O mRNA vaccine sera were able to efficiently neutralize the Omicron variant with titers ranging from 400 to 3200, however, none of them displayed neutralization effects on the WT strain or the Delta variant.

The present study compares side-by-side two mRNA-based SARS-CoV-2 vaccine candidates that express the RBD of the WT strain or the Omicron variant, respectively, for their immunogenicity and neutralization capacity. It was found that both of the mRNA vaccines are highly immunogenic, able to produce relatively high titers of neutralizing antibodies against the homologous strains. However, antibodies elicited by the two vaccines display distinct cross-neutralization profiles. Specifically, the RBD-WT mRNA vaccine sera exhibited slightly decreased (1.83-fold) neutralization potency on the Delta variant but were strikingly escaped by the Omicron variant (more than 22.6-fold reduction in neutralizing GMT), in consistent with the results obtained with human vaccinee sera^4,5,8^. Significantly, we found that the RBD-O mRNA vaccine sera potently neutralized the Omicron variant but failed to cross-neutralize the WT or Delta strains under the testing conditions used in this study, indicating a narrow neutralization spectrum for this vaccine. Based on the above results, we propose that a bivalent/multivalent vaccine formulation containing antigen components (such as RBD) derived from the Omicron variant and the WT strain (or an earlier variant such as Delta) should be developed in order to achieve broad protection against diverse SARS-CoV-2 strains/variants, especially Omicron and Delta.

In short, our current work demonstrates the feasibility and potency of an RBD-based mRNA vaccine specific for the SARS-CoV-2 Omicron variant, providing important information for further development of bivalent or multivalent SARS-CoV-2 vaccines with broad-spectrum efficacy.

## Acknowledgements

This work was supported by grants from the Chinese Academy of Sciences (XDB29040300) and from the Ministry of Science and Technology of China (2020YFC0845900). This project is also part of the European Union’s Horizon 2020 research and innovation program under grant agreement No.101003589. Chao Zhang is supported in part by Youth Innovation Promotion Association of the Chinese Academy of Sciences (CAS) and Shanghai Rising-Star Program (21QA1410000).

## Author contributions

Z.H. and H.K.W. conceived and designed the experiments; J-K.Z., C.Z. and Y.N.Y. participated in multiple experiments; Z.H., H.K.W., J.K.Z., C.Z. and Y.N.Y. analyzed the data; Z.H., H.K.W., J.K.Z., C.Z., and Y.N.Y. wrote the manuscript; Z.H., and H.K.W. provided the final approval of the paper.

## Conflict of interests

All authors declare that they have no competing interests.

## Notes

### Competing Interest Statement

The authors have declared no competing interest.

