## supplemental figure 1 for "An mRNA vaccine candidate for the SARS-CoV-2 Omicron variant"

### MATERIALS AND METHODS

#### Cells

HEK 293T cells were cultured in DMEM (Gibco, USA) supplemented with 10% fetal bovine serum (FBS; Gibco) at 37°C. HEK 293F cells were cultured in and FreeStyle 293 expression medium (Gibco). Human ACE2-overexpressing HEK 293T cell (293T-hACE2) were generated in a previous study[1].

#### Recombinant proteins and antibodies

RBD protein (residues R319 to S591) of SARS-CoV-2 strain Wuhan-Hu-1 (GenBank ID: MN908947.3) with a N-terminal strep-tag and C-terminal 6x His-tag was expressed in HEK 293F cells and then purified using Ni-NTA resin (Millipore).

A polyclonal antibody against HEK 293F-expressed RBD protein was generated in our previous study[1] and used for immunofluorescence staining analysis. A polyclonal antibody against *E. coli*-produced RBD protein was generated in our previous study[2] and used for western blotting analysis.

#### mRNA and LNP production

Construction of mRNA and LNP production was carried out according to a previously described protocol[3]. The codon-optimized WT or Omicron RBD genes (residues R319 to N532) with N-terminal T7 promotor, 5’-untranslated region (5’-UTR: 5’-AAATAAGAGAGAAAAGAAGAGTAAGAAGAAATATAAGAGCCACC-3’), interleukin-10 (IL-10) signal sequence, C-terminal 3’-UTR (5’-TGATAATAGGCTGGAGCCTCGGTGGCCATGCTTCTTGCCCCTTGGGCCTCCCCCCAGCCCCTCCTCCCCTTCCTGCACCCGTACCCCCGTGGTCTTTGAATAAAGTCTGA-3’) and poly-A tail of about 101 base pairs (bp) were synthesized and inserted into pUC57 vector. The firefly luciferase (FLuc) reporter gene with T7 promotor, 5’-UTR, IL-10 signal, 3’-UTR and poly-A tail was cloned into pUC57 vector, used as a control. The resulting plasmids were linearized and used to produce mRNAs *in vitro* using T7 RNA Transcription Enzyme Kit (Novoprotein; catalog No. E131-01A). The resultant mRNAs were purified and then capped using vaccinia capping system (New

England BioLabs, M2080S) and mRNA Cap 2´-O-Methyltransferase (New England BioLabs, M0366S) to produce the Cap I structure. The capped mRNAs were purified and stored at -80 °C until use.

LNP-mRNA formulations were prepared by rapid mixing as described previously[d,5]. Briefly, to generate LNP, D-Lin-MC3-DMA (MedChemExpress), DSPC (Avanti Polar Lipids), cholesterol (Sigma), and PEG-lipid (Avanti Polar Lipids) were solubilized in ethanol at a molar ratio of 50:10:38.5:1.5. The lipid mixture was added to 25 mM acetate buffer (pH 4.0) at a volume ratio of 1: 2 and incubated at room temperature for 2 min. Next, the lipid mixture was homogenized by extrusion through 100-nm pore-size filters (Genizer) using a lipid extruder (Genizer). For mRNA encapsulation, the capped mRNAs were solubilized in 25 mM acetate buffer and then mixed with LNP at a mRNA:lipid ratio of 0.056 mg/μmol, followed by incubation at 42 °C for 30 min. Formulations were dialyzed against PBS (pH 7.4) overnight, concentrated and passed through 0.45-μm filters before storage at 4 °C until use.

**Expression of the mRNA in HEK 293T cells**

*In vitro* transcribed mRNAs were transfected into HEK 293T cells in 6-well or 24-well plates using Lipofectamine 2000 Transfection Reagent (Invitrogen) following manufacturer's instructions. At 48 h post-transfection, the culture supernatants were collected and concentrated for western blot analysis with HRP-conjugated anti-His tag antibody (Proteintech) and anti-RBD polyclonal antibody as detection antibodies. For immunofluorescence staining analysis, at 48 h post-transfection, the cells were washed with PBS, fixed with 4% paraformaldehyde and permeabilized, followed by blocking with 10% FBS and 10% BSA in PBS for 1 h. The fixed cells were incubated with mouse anti-RBD polyclonal antibody, followed by Alexa-488-conjugated anti-mouse secondary antibody (Invitrogen). Nuclei were stained with 4,6-diamidino-2-phenylindole (DAPI) in PBS. Finally, the stained cells were observed under a fluorescence microscope (Olympus).

**Animal experiments**

All the animal experiments in this study were approved by the Institutional Animal Care and Use Committee at the Institut Pasteur of Shanghai.

For bioluminescence imaging to detect *in vivo* distribution of Fluc-mRNA-LNP, female BALB/c mice aged 6-8 weeks (n = 5) were injected with 10 μg of the FLuc-mRNA-LNP via intramuscular route. At 6-, 12-, 24-, and 48-h after the mRNA-LNP injection, the animals were injected intraperitoneally (i.p.) with luciferase substrate (Promega). After reaction for 8 minutes, fluorescence signals were recorded by IVIS Spectrum instrument (PerkinElmer) and analyzed using Living Image software 3.0.

For animal immunization, groups of adult female BALB/c mice (8 mice per group) were intramuscularly immunized with the LNP-encapsulated RBD-WT or RBD-O mRNA vaccines (10 μg of mRNA per dose) at weeks 0 and 2. Another group of mice (5 mice per group) were injected with the FLuc-mRNA-LNP formulation (10 μg of mRNA per dose), used as a control. Blood samples were collected from individual mice at weeks 2 and 4 for antibody analysis.

**Enzyme-linked immunosorbent assay (ELISA)**

ELISA were carried out for evaluation of antigen-specific antibodies. Briefly, 96-well plates were coated with 25 ng/well of HEK 293F-expressed RBD protein at 4 °C overnight. After blocking with 5% milk, individual antisera were diluted 1:100 and added to the plates, followed by incubation for 2 h at 37 °C. The plates were incubated with HRP-conjugated anti-mouse IgG (Sigma) for 1 h at 37 °C. After color development, absorbance at 450 nm was measured.

**Pseudovirus neutralization assay**

Murine leukemia virus (MLV)-based pseudoviruses bearing spike proteins of WT (Wuhan-Hu-1 strain), Delta, or Omicron SARS-CoV-2 strains were produced according to our previously reported protocol [2]. Mutations in the Delta S protein include T19R, del156-157, R158G, L452R, T478K, D614G, P681R and D950N. Mutations in the Omicron S protein include A67V, H69delV70del, T95I, G142D-V143del-Y144del-Y145del, N211del-L212I, ins214EPE, G339D, S371L, S373P, S375F, K417N, N440K, G446S, S477N, T478K, E484A, Q493R, G496S, Q498R, N501Y, Y505H, T547K, D614G, H655Y, N679K, P681H, N764K, D796Y, N856K, Q954H, N969K, and L981F.

All sera samples were 2-fold serially diluted and subjected to pseudovirus neutralization assay as previously described [2]. The luciferase activity was measured using the luciferase assay system (Promega). 50% pseudovirus neutralization titers (NT50) were calculated using nonlinear regression in GraphPad Prism (version 8).

***Statistics analysis***

All statistical analyses were performed using GraphPad Prism software v6.0. Statistical significance was analyzed using Student's *t*-test.

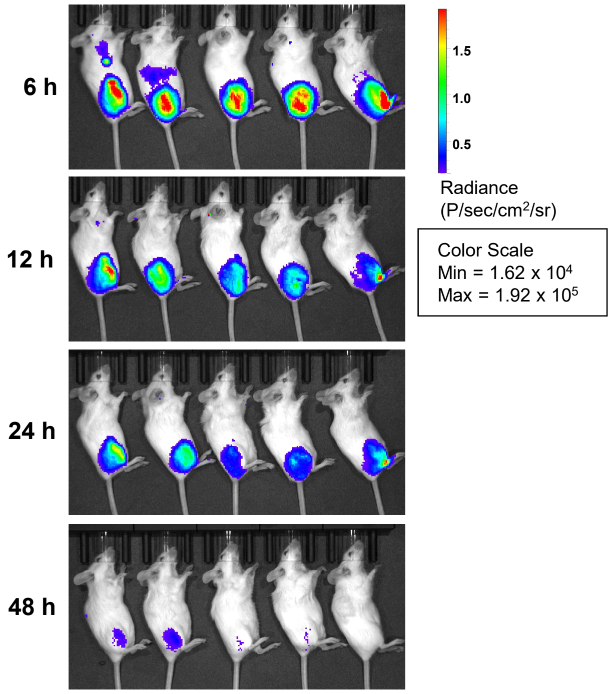


**Supplementary Fig. S1** Characterization of the mRNA-LNP expression in vivo. Mice were injected with the FLuc-mRNA-LNP formulation and then subjected to bioluminescence imaging at different time points.
